# An efficient and scalable pipeline for epitope tagging in mammalian stem cells using Cas9 ribonucleoprotein

**DOI:** 10.1101/255737

**Authors:** Pooran Singh Dewari, Benjamin Southgate, Katrina Mccarten, German Monogarov, Eoghan O’Duibhir, Niall Quinn, Ashley Tyrer, Colin Plumb, Carla Blin, Rebecca Finch, Raul Bardini Bressan, Gillian Morrison, Ashley M. Jacobi, Mark A. Behlke, Alex von Kriegsheim, Simon Tomlinson, Jeroen Krijgsveld, Steven M. Pollard

## Abstract

CRISPR/Cas9 can be used for precise genetic knock-in of epitope tags into endogenous genes, simplifying experimental analysis of protein function. However, Cas9-assisted epitope tagging in primary mammalian cell cultures is often inefficient and reliant on plasmid-based selection strategies. Here we demonstrate improved knock-in efficiencies of diverse tags (V5, 3XFLAG, Myc, HA) using co-delivery of Cas9 protein pre-complexed with two-part synthetic modified RNAs (annealed crRNA:tracrRNA) and single-stranded oligodeoxynucleotide (ssODN) repair templates. Knock-in efficiencies of ~5-30%, were achieved without selection in embryonic stem (ES) cells, neural stem (NS) cells, and brain tumour-derived stem cells. Biallelic-tagged clonal lines were readily derived and used to define Olig2 chromatin-bound interacting partners. Using our novel web-based design tool, we established a 96-well format pipeline that enabled V5-tagging of sixty different transcription factors. This efficient, selection-free and scalable epitope tagging pipeline enables systematic surveys of protein expression levels, subcellular localization, and interactors across diverse mammalian stem cells.

## Introduction

Defining the protein levels, subcellular localisation and biochemical interactions for the >20,000 protein-coding genes in the mammalian genome remains a formidable challenge. Ideally these would be explored across diverse primary cells – rather than genetically transformed and corrupted cell lines. Large-scale projects, such as the human protein atlas, have performed systematic characterisation of antibodies against native proteins (Thul et al. 2017). However, there are inherent difficulties in discovering, validating and distributing high-quality antibodies that cover all key applications and species (e.g. immunoblotting, immunoprecipitation, ChIP-Seq and immunocytochemistry).

A complementary strategy is to use epitope tagging: the fusion of small peptide-coding sequences with to a protein of interest (e.g. 3XFLAG, HA, V5, and Myc) (Jarvik and Telmer 1998). In contrast to plasmid-driven cDNA overexpression, which creates artificially high protein levels, the knock-in of small epitope tags to endogenous genes provides physiologically relevant levels. A small set of pre-validated tag-specific antibodies is then used across diverse downstream experimental applications. This has been a key tool in yeast studies, resulting in global characterization of core protein complexes and their extensive interaction networks (Gavin et al. 2006; Krogan et al. 2006). However, to date this approach has not been widely adopted in mammalian cells, primarily due to the poor efficiency of homologous recombination (HR). Repurposed programmable nucleases now provide a potential solution.

Formation of a site-specific DNA double-strand break (DSBs), will massively enhance HR-mediated repair. This has become straightforward with the discovery and application of clustered regularly interspaced short palindromic repeats (CRISPR) and CRISPR-associated (Cas) proteins as designer site-specific nucleases. CRISPR/Cas9 was uncovered as a form of microbial adaptive immunity (Bhaya et al. 2011) that has been repurposed for genome editing in mammalian cells (Cong et al. 2013; Doudna and Charpentier 2014).

Cas9 is an RNA-guided endonuclease that binds complementary DNA via formation of a 20 bp RNA:DNA heteroduplex. Stable binding of Cas9/gRNA complex at the target site leads to activation of nuclease domains and formation of a double-stranded DNA break (DSB) (Jinek et al. 2014). DSBs are predominantly repaired through the error-prone non-homologous end joining (NHEJ) pathway, resulting in insertion or deletion mutations (indels) (Sander and Joung 2014). Alternatively, at lower frequencies HR-mediated repair will occur.

Successful knock-in of tags therefore requires co-delivery of three ingredients to the mammalian cell: the Cas9 protein, a guide RNA, and a donor repair template (i.e. single or double-stranded DNA encoding the tag or reporter with homology arms). These CRISPR components are typically delivered via transient plasmid transfection or viral vectors. Bespoke targeting vector plasmids are usually constructed for delivery of large cargoes or conditional alleles. Selection strategies, such as flow cytometry, or use of antibiotic resistance cassettes, are then used to enrich for edited cells.

Despite the successes of current approaches (Dalvai et al. 2015; Savic et al. 2015; Mikuni et al. 2016; Xiong et al. 2017), it is clear many bottlenecks restrict widespread applications; i) Production of tailored plasmids for each component can be tedious and time-consuming; ii) Plasmid-based selection strategies often leave ‘scars’ in the genome that might interfere with normal regulatory processes; iii) Plasmid DNA, either Cas9/gRNA expression vectors or targeting vectors, can randomly integrate in the genome causing insertional mutagenesis or increasing risks of off-target cleavage; iv) current strategies are often not readily scalable to enable routine exploration of large numbers of genes; v) recovery of biallelic clonal lines is inefficient. Thus, there remains an unmet need for improved and knock-in strategies that can work efficiently in primary mammalian cells, such as pluripotent (ES/iPS), multipotent (tissue stem cells) and cancer stem cells.

Improved efficiencies of CRISPR editing in mammalian cells have been demonstrated using recombinant Cas9 protein (Kim et al. 2014; Ramakrishna et al. 2014; Bressan et al. 2017). Cas9 protein is complexed with in vitro transcribed (IVT) RNA, to produce a ribonucleoprotein (RNP) that can be then delivered into cells. The RNPs are short-lived and cleared by cells within 24-48 hours, reducing the risk of both formation of mosaic clones or off-target cleavage (Kim et al. 2014; Lin et al. 2014; Zuris et al. 2015; Cameron et al. 2017). However, RNP-assisted methods have mainly been used for gene knock-out (Kim et al. 2014; Liang et al. 2015), incorporation of point mutations (Ma et al. 2017; Rivera-Torres et al. 2017), or knock-in of restriction enzyme sites (Lin et al. 2014; Schumann et al. 2015). We and others have recently made use of RNP with in vitro transcribed (IVT) sgRNAs for CRIPSR epitope tagging (Bressan et al. 2017; Liang et al. 2017), yet the efficiencies in the absence of selection are highly variable, and this approach cannot be scaled.

To avoid IVT, chemically-modified ~100 nt-long sgRNAs can be synthesized (Hendel et al. 2015); however, these are prohibitively expensive, limiting applications. Alternatively, a two-part chemically synthesized short target-specific crRNA plus longer generic tracrRNA, can be used. This is cheaper (only the crRNA needs to be resynthesized for new targets) and has better performance (Aida et al. 2015; Anderson et al. 2015). This ‘dual-RNA’ approach has been advanced further by ‘base, backbone and end’ modifications of cr/tracrRNAs (Rahdar et al. 2015; Kelley et al. 2016) and use of shorter and more effective modified cr/tracrRNAs (Jacobi et al. 2017). Modified synthetic cr/tracrRNAs are resistant to nuclease digestion, limit cellular immune responses, have greater stability, and therefore provide enhanced targeting efficiency (Jacobi et al. 2017).

Here we explored whether RNPs with synthetic modified two-part guide RNA can enhance efficiency of epitope tagging in primary mammalian stem cells. We find that co-delivery of a Cas9 RNP (dual-synthetic RNA) with ~200 bp single-stranded oligodeoxynucleotides (ssODNs) supports highly efficient epitope tagging. This is achieved across a variety of stem cell types without any requirement for plasmids, selection steps, flow cytometry-based enrichment, or IVT reactions. To provide proof or principle, we developed a novel web-based design tool and demonstrate effective tagging in 96 well-plate format. We demonstrate one application, by identifying Olig2 protein partners using immunoprecipitation-mass spectrometry (IP-MS).

## Results

### Cas9 protein complexed with synthetic cr/tracr RNAs enables highly efficient epitope tagging in neural and glioma stem cells

sgRNAs produced by IVT reactions can vary in quality and quantity and are prone to degradation, either during production and/or following delivery into cells. We therefore explored a synthetic modified two-part guide RNA system (annealed 36-mer crRNA: 67-mer tracrRNA) (Anderson et al. 2015; Jacobi et al. 2017).

Guide RNAs were designed to cut proximal to the stop codon in the 3’ UTR of *Olig2* or *Sox2* (Figure 1A). The efficacy of custom synthetic modified RNAs (csRNAs) was compared to IVT-generated sgRNAs. RNA was complexed with recombinant Cas9 protein and transfected into a primary adult mouse neural stem (NS) cells (ANS4), using an optimised nucleofection program. RNP was delivered together with a ~200bp single-stranded DNA donor encoding the V5 tag, flanked with ~70 nucleotide homology arms (Figure 1B). After 5 days, cells were analysed using immunocytochemistry (ICC) for the V5 protein fusion (Figure 1C). The csRNA-based RNP gave a >4-fold and >10-fold increase in knock-in efficiency for *Olig2* and *Sox2*, respectively (Figure 1D). Improved knock-in efficiencies were also obtained using an independent cell line (glioma initiating neural stem cells; termed IENS) (Figure 1D). PCR genotyping and Sanger sequencing confirmed in-frame and error-free insertion of V5 tag sequence at the C-terminus of *Olig2* and *Sox2* loci (Supplementary Figure 1A). V5 positive cells all displayed the anticipated nuclear localization and levels with no indication of non-specific expression.

**Figure 1.**
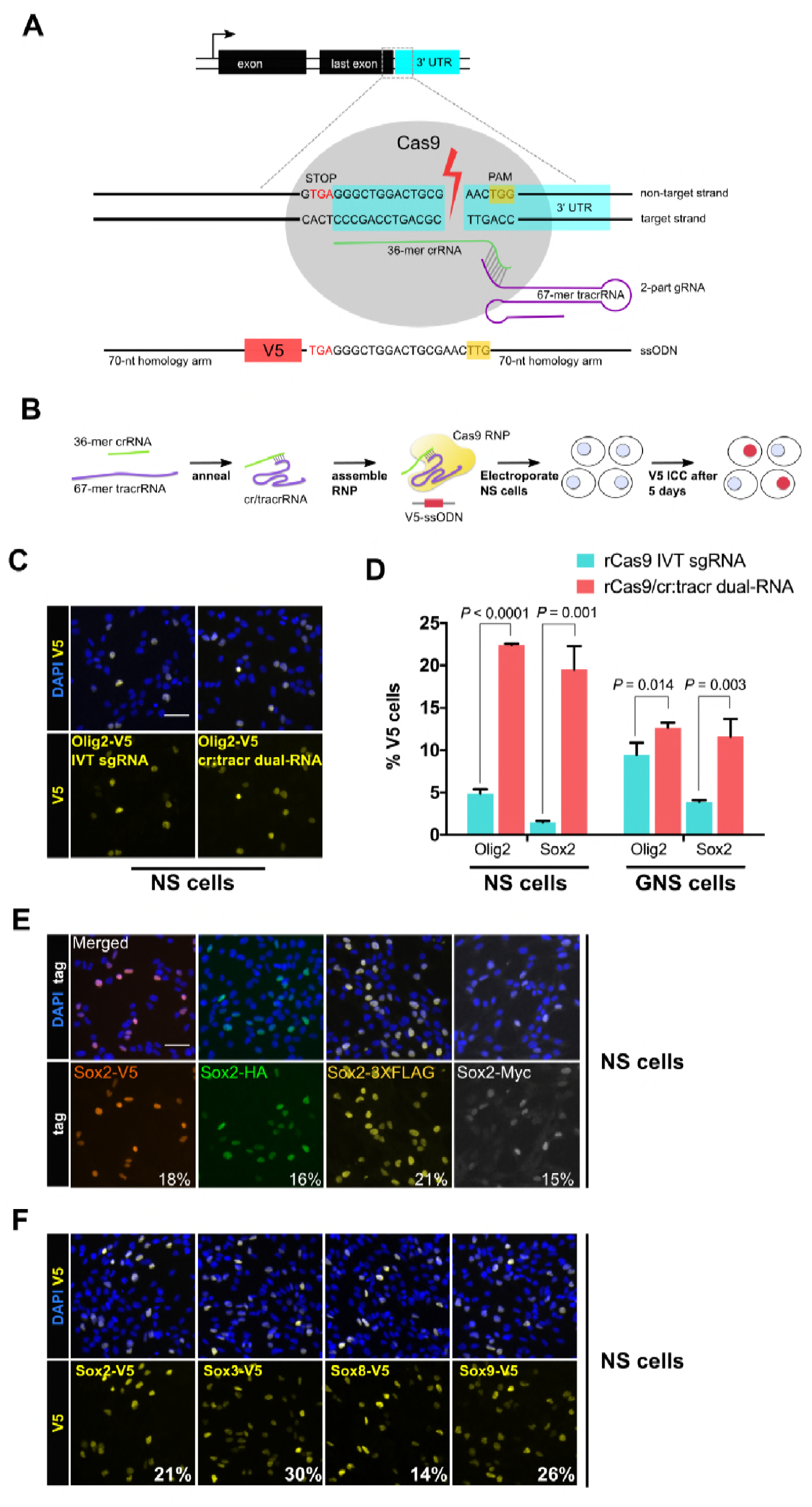
Cas9 protein in complex with synthetic cr/tracrRNAs enables highly efficient knock-in of biochemical tags in mouse neural and glioma stem cells. **(a)** Schematic representation of epitope knock-in strategy. A crRNA was designed against the 3’UTR of each target gene. Target site with double-stranded break is shown with Cas9 RNP (grey), PAM in yellow box, and single-stranded donor DNA that harbours PAM-blocking mutations and V5 tag coding sequence flanked by 70-mer homology arms on both sides. **(b)** Cas9 RNP complexes were assembled *in vitro* by incubation of recombinant Cas9 protein with either IVT sgRNA or synthetic two-part cr:tracrRNA and electroporated into NS cells. V5 ICC was to quantify knock-in. **(c)** Representative ICC images for the detection of Olig2-V5 fusion protein in the bulk populations of transfected cells. **(d)** HDR-mediated insertion of V5 tag was determined by scoring V5-positive cells (%) in the bulk populations of transfected cells. Results from three independent experiments are shown for *Sox2* and *Olig2* V5 tagging using mouse neural stem (NS) and glioma neural stem (GNS) cells. Error bars indicate standard deviation values based on a minimum of two experiments, P-values were derived using unpaired t test. **(e)** ICC for *Sox2* gene epitope tagging at the C-terminus with V5, HA, 3XFLAG, or Myc epitope. Numbers represent percentage of tagged cells in the bulk population for each tagging experiment. **(f)** Representative bulk population V5 ICC images for Sox2, Sox3, Sox8, and Sox9 V5 knock-in are shown. Average knock-in efficiency from two independent experiments is shown at the bottom (numbers in white).

Knock-in efficiency might vary when using distinct biochemical tags. We therefore tested a variety of widely used alternative tags (V5, HA, 3XFLAG, or Myc). Each tag varied substantially in size, and consequently homology arm. Nevertheless, we observed similar rates of knock-in (15%-21%) across all four tags for *Sox2* (Figure 1E and Supplementary Table 1). An independent adult NS cell culture also gave similar results (9%-15% knock-in efficiency, Supplementary figure 1D).

High knock-in efficiencies were not limited to *Sox2* and *Olig2*. We found that *Sox3*, *Sox8*, and *Sox9* – three *Sox* family members expressed in NS cells – had knock-in efficiencies of 30%, 14% and 26%, respectively (Figure 1F; Supplementary Figure 1B and 1C). Furthermore, we could simultaneously knock-in two different tags (*Sox2*: HA tag; and *Olig2*: V5 tag) in the same cells using a single transfection (4% double-positive cells) (Supplementary Figure 1E).

Altogether, these results indicate that use of the modified 2-part synthetic cr/tracrRNA system is more effective than IVT for epitope knock-in in mammalian NS and GNS cells. Using csRNA-based RNP (csRNP), we achieved 5-30% knock-in efficiency across distinct cell lines for different genes and using different tags. This was accomplished without the requirement for flow cytometry or plasmid-based selection strategies.

### The high frequency of knock-in using csRNP facilitates simple recovery of biallelic-tagged clonal lines

Generation of biallelic-tagged clonal lines is advantageous for downstream applications, as all target protein will be tagged, enabling improved signal to noise ratios in assays. This gives greater confidence in the results obtained. Low efficiency of tagging provides means that many hundreds or thousands of clones need to be screened and genotyped, limiting downstream applications. Heterozygous lines might also harbour a indels on the other, non-HR tagged, allele (Merkle et al. 2015; Bressan et al. 2017); inappropriate transcriptional/post-transcriptional regulation may result.

The improved knock-in efficiency using the csRNP method encouraged us that recovery of biallelic clonal lines might be straightforward. Clonal NS cell lines were established from bulk tagged populations following single-cell deposition into to 96-well plates. Tagged clones were then identified following replica plating and ICC for V5 (Figure 2A). Biallelic clones were scored using PCR primer-pairs flanking the tag sequence (Figure 2B) and validated using ICC and Sanger sequencing (Figure 2C and Supplementary Figure 2).

**Figure 2.**
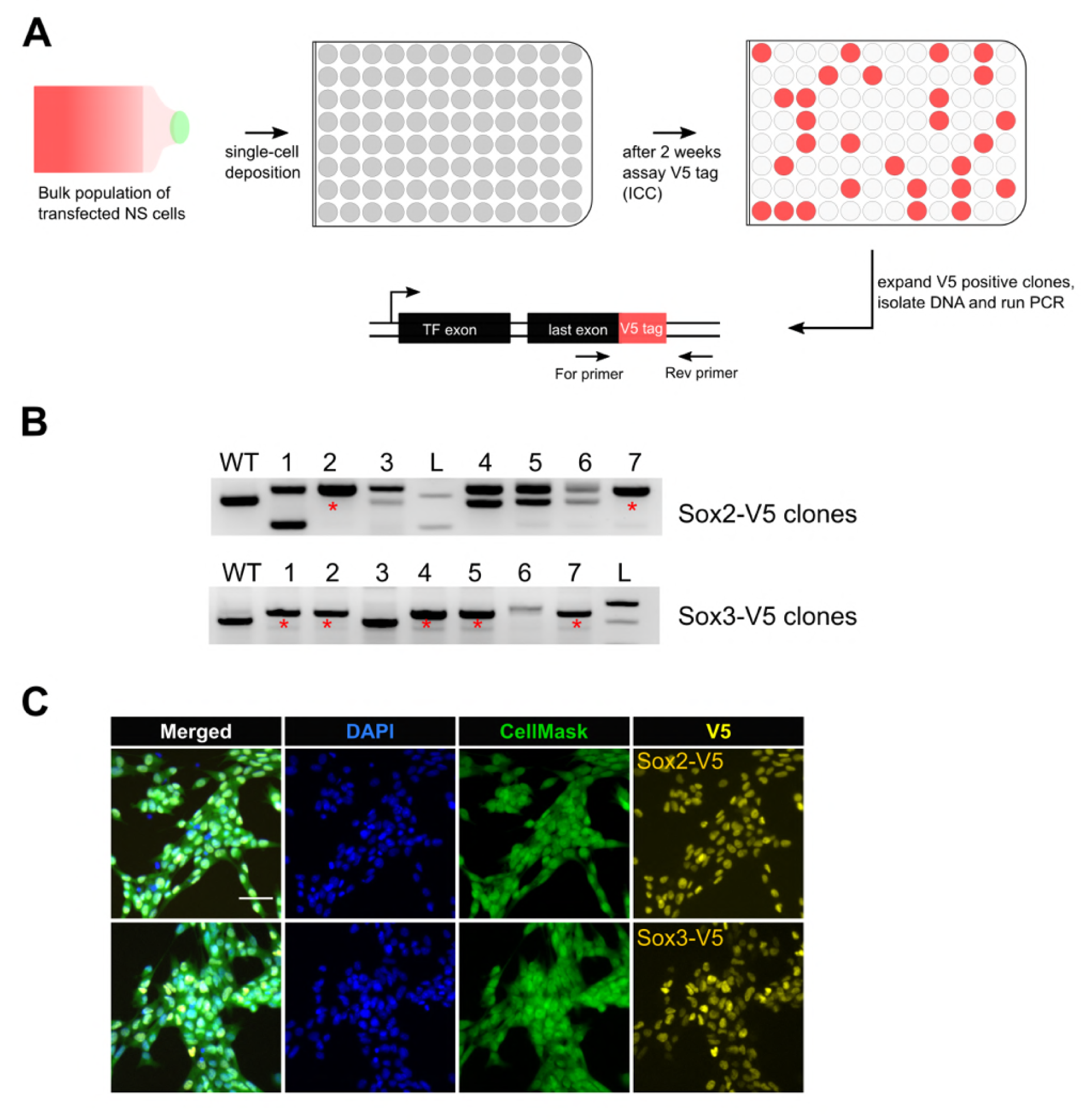
Bi-allelic knock-in clonal lines can be readily generated. **(a)** Experimental workflow for generating clonal lines. Cells from the bulk population cultures were single-cell deposited to 96-well plates using flow cytometry. Two weeks later, V5 positive clones were selected by ICC assay and confirmed by PCR genotyping. **(b)** Representative agarose gel pictures for Sox2-V5 and Sox3-V5 PCR genotyping are shown. Wild-type unedited cells (control) are marked with WT; 1kb+ DNA ladder is marked with L; bi-allelic clones are marked with red asterisks. **(c)** ICC images of bi-allelic V5 knock-in clones are shown for Sox2 and Sox3 TFs. Nuclear stain DAPI (blue), entire cell stain CellMask^TM^ HCS (green), and V5 tag (yellow) are shown.

89 clonal lines were generated from the *Sox2*-V5 knock-in cells. 13 of these were V5-positive by ICC. Of these four had successfully integrated the V5 tag sequence at both alleles (30% of V5-positive clones) (Table 2). High frequencies of bi-allelic knock-in were also obtained for *Sox3*-V5 (62.5%). We also derived several biallelic knock-in lines from another independent cell line (IENS, mouse glioma NS cells): *Sox2, Sox3, Sox8* and *Sox9* (7%, 26%, 57%, and 15% biallelic knock-in clones, respectively) (Table 2). Thus, biallelic tagged clonal lines can be readily recovered.

### Multiple stem cell-types can be epitope-tagged using csRNPs, including non-expressed genes

To test the general applicability of the csRNP tagging method across other types of stem cells, we compared head-to-head tagging efficiencies between mouse ES cells and NS cells. We initially focussed on four transcription factors (TFs): *Sox2, Sox3, Ctcf* and *Pou3f1*, and the chromatin regulator *Ezh2;* each of these is expressed in both cell types. In each case we found that mouse ES cells (E14Tg2a) were tagged at similar level of efficiency to the NS cells (Figure 3A and 3B).

**Figure 3.**
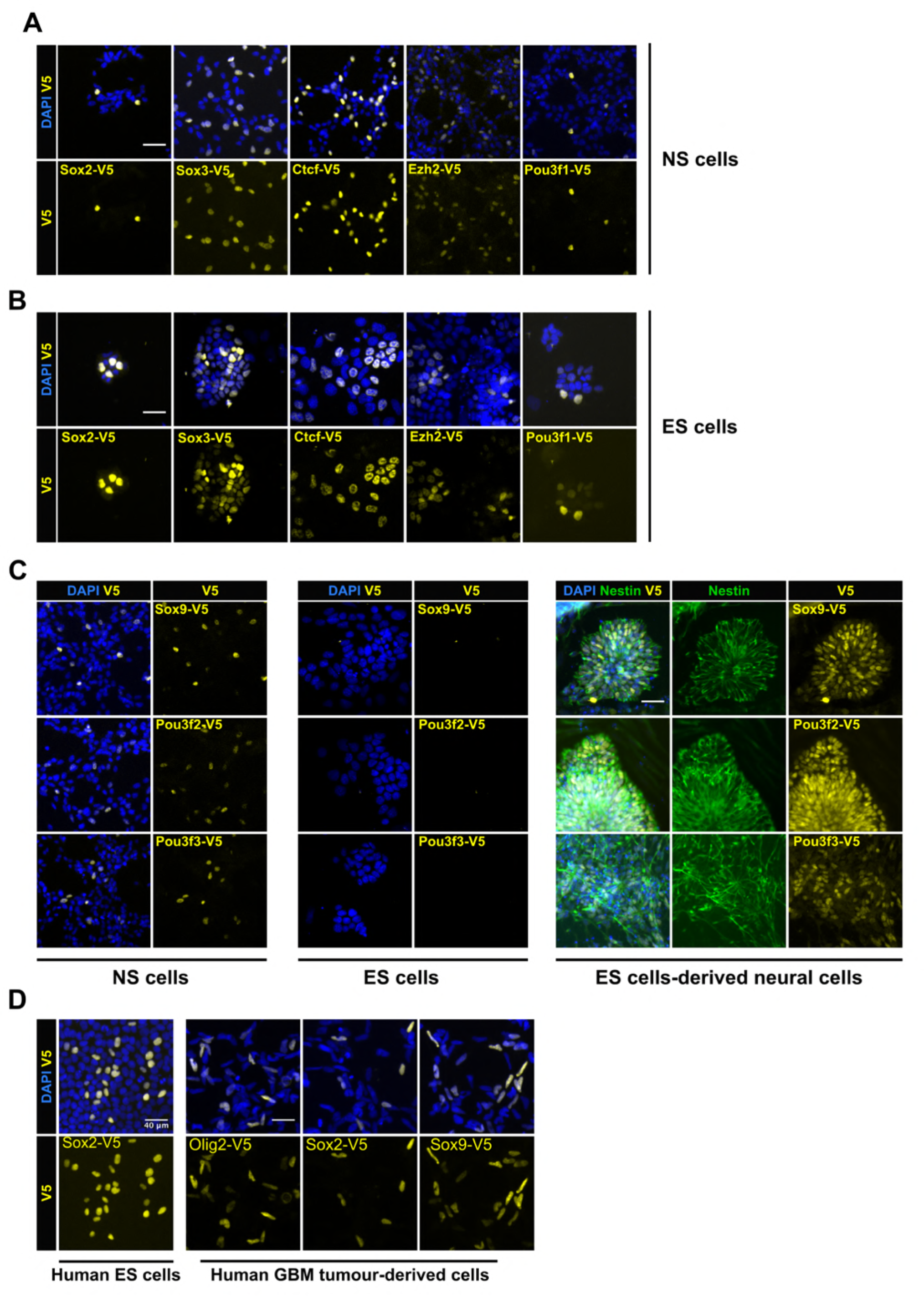
Variety of cell types can be epitope-tagged using RNP method. Representative V5 ICC images are shown for V5 tagging of five TFs that are expressed in both mouse NS cells **(a)** and ES cells **(b)**. V5 ICC images for the three neural-specific TFs Sox9, Pou3f2, and Pou3f3 in mouse NS cells **(figure c, left panel)**. Mouse ES cells were electroporated with similar three TF-reagents **(figure c, middle panel)**. Later, ES bulk populations of non-expressed TFs from were differentiated into neural lineage and assayed for expression of V5 fusion proteins by ICC **(figure c, right panel)**, differentiation into neural stem progenitors was confirmed by Nestin ICC (in green). **(d)** V5 ICC images for epitope tagging using human ES cells and GBM patient-derived cell lines.

Non-expressed genes are often difficult to engineer. We therefore tested csRNPs for several neural-affiliated TFs (*Sox9, Pou3f2,* and *Pou3f3*) that are expressed in NS cells but not ES cells (Figure 3C). V5 insertion was first confirmed by PCR genotyping in the bulk populations and suggested both ES cells and NS cells were effectively tagged (Supplementary Figure 3). V5-tagged protein was detected by ICC only in NS cells and not in ES cells (Figure 3C, left and middle panels). However, for each of these genes, upon neural lineage differentiation of the ES cells we noted Nestin-expressing neural rosettes that were V5-positive (Figure 3C, right panel). Thus, non-expressed genes can be successfully tagged in ES cells, without deploying any selection strategies or plasmids. We also assessed tagging in human ES cells. V5 knock-in was successfully demonstrated in human ES cells for *SOX2* (Figure 3D). We also tested human GBM-derived primary cells for *OLIG2*, *SOX2*, and *SOX9* genes (15%-70% efficiency) (Figure 3D).

These data illustrate the power of the efficient tagging to knock-in of non-expressed genes in stem cells, and subsequent monitoring of the tagged protein in their differentiating progeny. We also conclude that the same csRNP epitope tagging approach and reagents can works effectively across diverse mouse and human pluripotent stem cells, neural stem cells and cancer-derived stem cells.

### DSB distance from the stop codon is a key parameter for successful tagging

To further define the parameters influencing the reliability and efficiency of tagging, we attempted V5 epitope knock-in at the C-terminus for all 50 *Sox* and *Fox* genes. This set of genes included both expressed and non-expressed genes. Previous studies have indicated that the distance of the DSB to the insertion site influences the frequency of successful HR (Bialk et al. 2015; Paquet et al. 2016). We designed two different target crRNAs in the 3’UTR of the gene for each of the 50 target genes; one cutting proximal and the other distal to the stop codon. Cells were transfected with Cas9 RNP containing either of the crRNA and ssODN to assess if distance of the cut site from the stop codon influenced knock-in efficiencies (Figure 4A).

**Figure 4.**
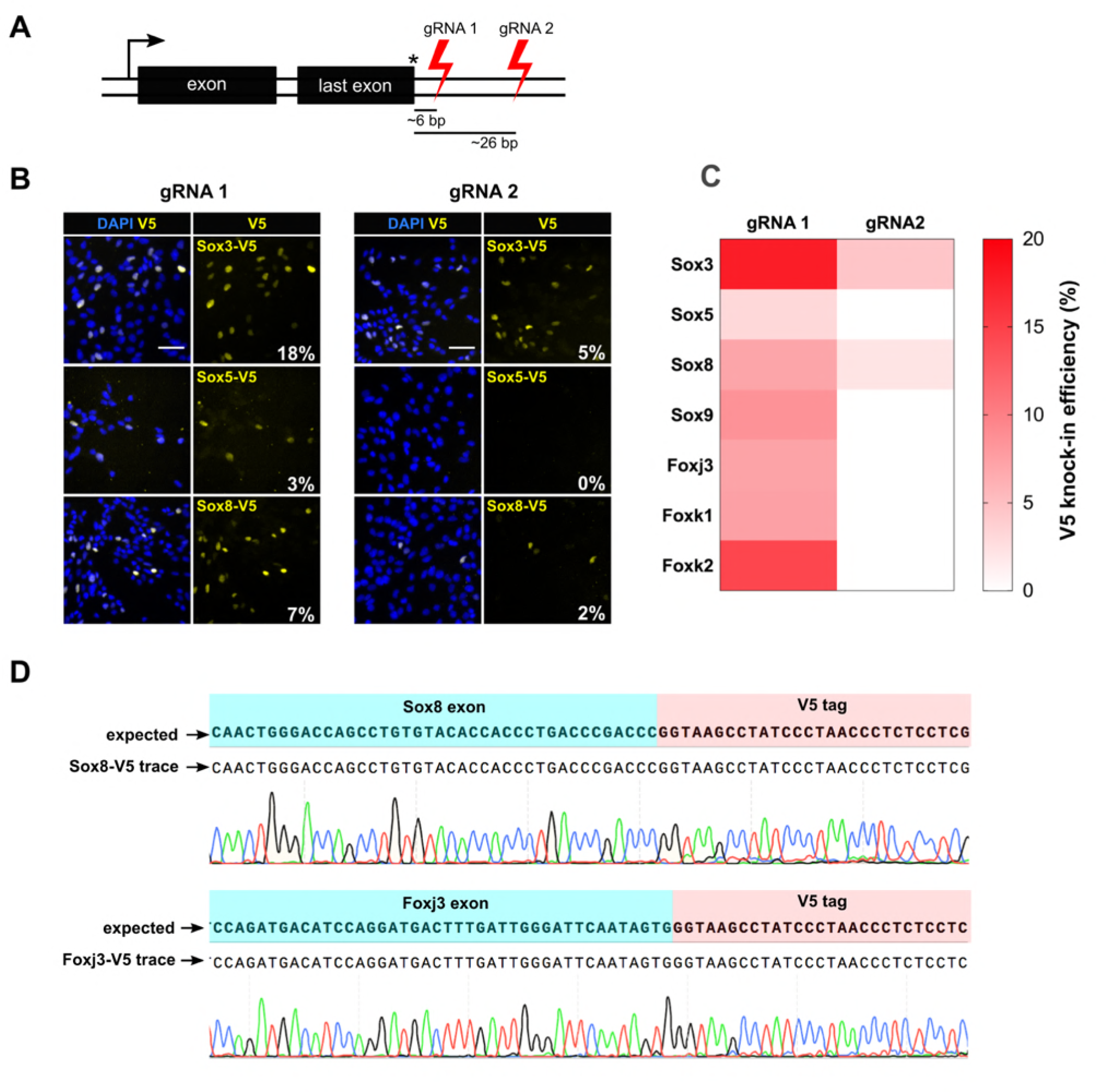
RNP cutting proximal to insertion site is more efficient and allows knock-in of non-expressed genes. **(a)** Schematic showing the relative position of guide RNAs in the 3’ UTR. The first set of gRNA (gRNA1) cuts proximal to the stop codon, the second set gRNA 2 distal. Average distance of cut site from the insertion site is shown for both sets. **(b)** Representative V5 ICC images of Sox3, Sox5, and Sox8 V5 knock-in bulk populations obtained with gRNA 1 (left panels) and gRNA 2 (right panels) are shown; % V5 knock-in efficiency for each experiment is shown at the right bottom of V5 panels (numbers in white). **(c)** Heatmap representation of V5 knock-in efficiency obtained with gRNA1 and gRNA2 for different genes, color code is shown on the right. **(d)** Example of sequencing traces using the bulk populations of V5-tagged cells for *Sox8* and *Foxj3*. Alignment with the expected TF-V5 fusion protein is shown.

By PCR genotyping we found that both proximal and distal gRNAs could result in successful tagging in the majority of cases; 30/50 genes (60%) for proximal DSB and 27/50 for the distal DSB (54%) (Supplementary Figure 4 and 5). However, this assay is qualitative. To quantitatively score the knock-in efficiency, we performed V5 ICC assay for the seven expressed TFs (Figure 4B and C). Sanger sequencing confirmed targeted insertion of the V5 tag-coding sequence (Figure 4D). Importantly, by comparing the efficiency of tagging for these 7 genes, we noted a consistent trend towards increased tagging for the most proximal cut site. For example, Sox3 showed 18% and 5% knock-in efficiency, for proximal and distal gRNAs, respectively (Figure 4B). For four genes (*Sox9, Foxj3, Fok1* and *Foxk2*), the distal gRNA did not work, whereas the proximal worked well (Figure 4C). There were no genes for which the more distal gRNA worked better than the proximal gRNA. These results suggest proximity of DSB to the stop codon influences the efficiency of knock-in.

### A scalable pipeline for high-throughput knock-in of epitope tags using 96-well microplates

It is often desirable to explore large numbers of proteins within a shared family, complex or pathway. Methods enabling knock-in of many genes in parallel would be valuable. As the gene-specific synthetic short crRNAs and matched ssODN repair templates can be obtained from commercial suppliers in 96-well microplates. Indeed, all steps, can be performed easily in 96 well format: preparation of the transfection-ready components via automated liquid handling, benchtop incubation/annealing, 96-well transfection, and automated microscopy to acquired images across 96 well plates. We reasoned that scale-up could therefore be relatively straightforward. A major remaining bottleneck, however, is the need for bioinformatics design tools specifically tailored towards epitope tagging applications.

Manually extracting gene sequence data, identifying appropriate gRNAs, and design of modified repair ssODNs, would be laborious and error-prone for hundreds of genes. To automate the design, we developed a novel web-based tool. This enables design of appropriate gRNA and ssODNs for both human and mouse species, with flexibility in choice of tag. Our ‘Tag-IN’ incorporates key design rule and parameters – e.g. incorporating “Rule Set 2” for maximised activity (Doench et al. 2016), and “MIT” scoring model (Hsu et al. 2013) to minimize off-target effects. Our tool also considers optimal distance from the insertion site (stop codon), and outputs the matched ssODNs modified with PAM-blocking mutations and appropriate epitope tag sequence. ‘Tag-IN’ also enable batch design; critical for the effective scale up to 96 well format (Figure 5).

**Figure 5.**
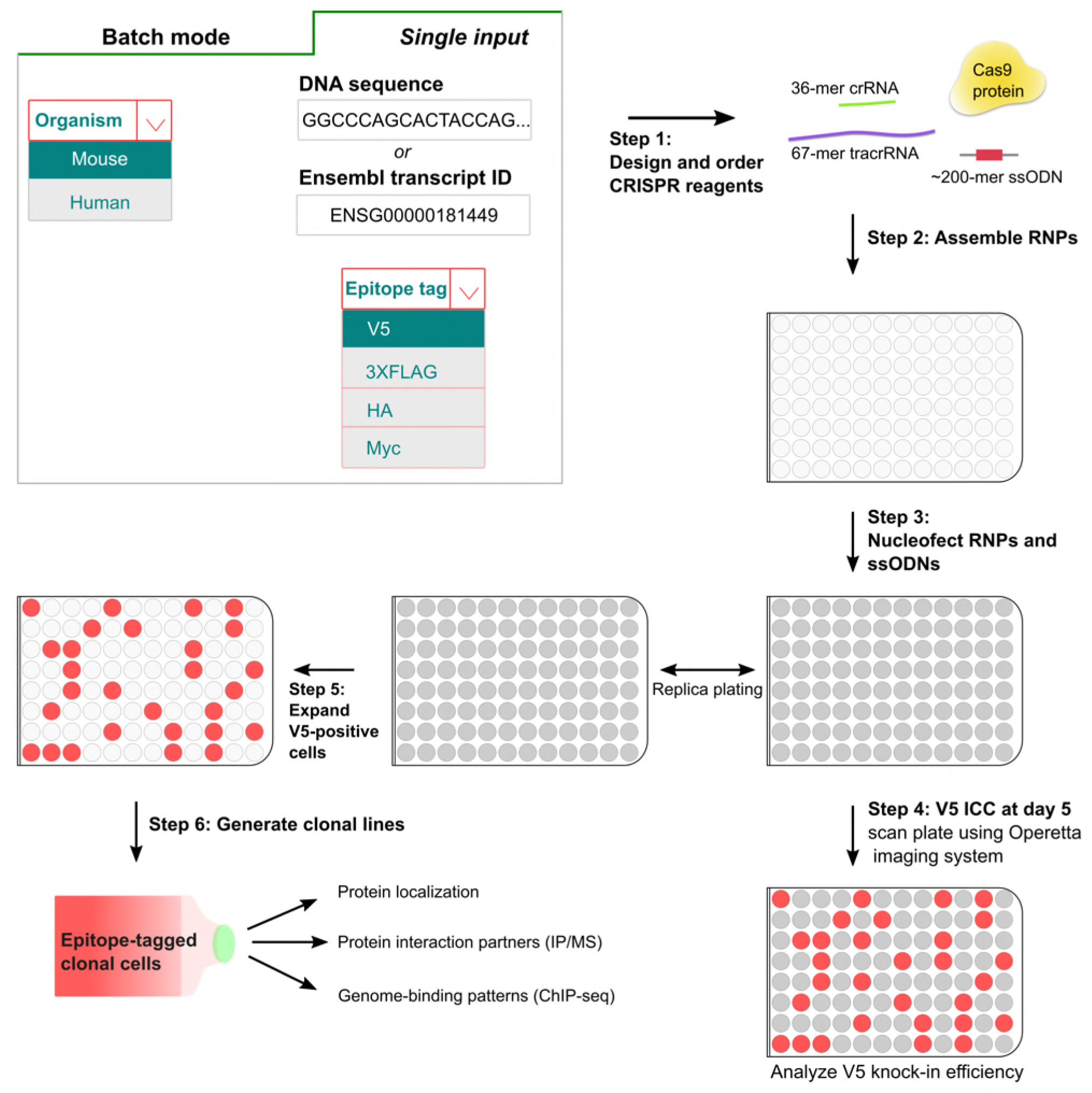
A simple pipeline for high-throughput epitope knock-in using 96-well format. In the first step, crRNAs and matched ssODNs are designed (either single input or batch processing) using the ‘Tag-IN’ bioinformatics tool. The tool picks top two crRNAs in the 3’UTR (within 8-15 bp from stop codon if high-quality crRNAs available) and designs an ssODN for each query based on user’s choice of the epitope tag. After the procurement of CRISPR ingredients from the supplier, the RNPs and matched ssODNs are assembled in vitro in 96-well plates (step 2) and transfected into stem cells using Amaxa shuttle system (step 3). Five days after the transfection, a replica plate is processed for V5 ICC and images are captured using high-content imaging system Operetta (step 4). V5-positive cells from the corresponding wells can be later expanded to derive clonal lines for downstream applications (step 5 and 6).

Using the Tag-IN tool, we designed crRNAs and matched V5 encoding repair ssODNs against 185 different transcription factors. These genes were selected based on expression in human glioblastoma stem cells. One gRNA was tested for each gene. The RNPs were prepared using a 96-well head liquid handling device (Felix, CyBio) and then transfected in parallel into mouse GNS cells (Figure 5). Five days later, ICC was performed and V5 knock-in efficiencies quantified across the entire plate using automated plate image capture (Operetta high-content imaging system, Perkin Elmer). *Sox2* tagging was used as a positive control in six wells; these gave consistent V5 knock-in efficiency across the plate, confirming no edge effects during the procedure (10.5 ± 2.5%) (Figure 6A).

**Figure 6.**
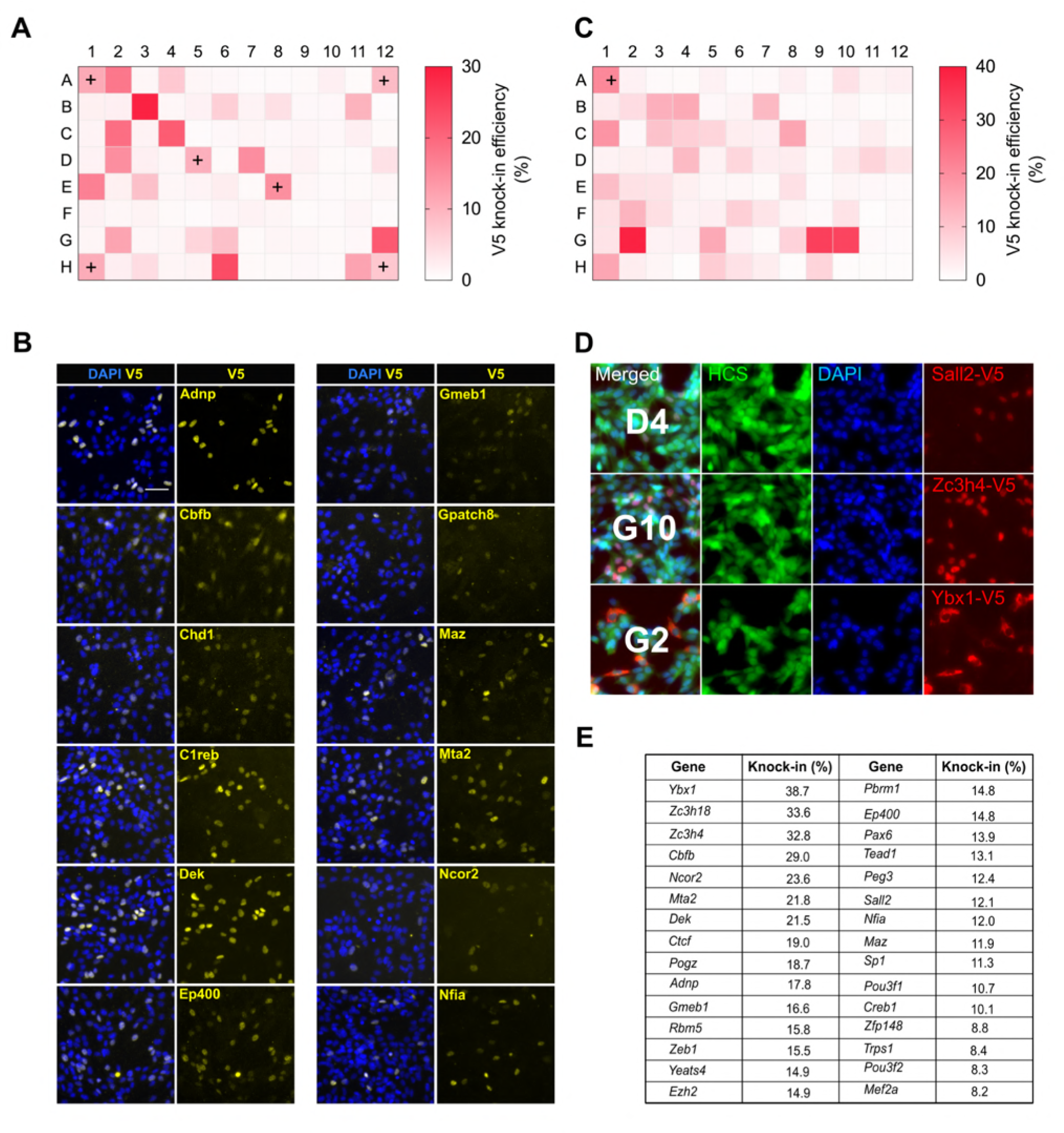
High-throughput epitope knock-in in a 96-well format. (**a**) Heat map of V5 knock-in efficiency across the entire 96-well plate. The positive control Sox2 was included in 6 different wells (marked with ‘+’). (**b**) Representative ICC images of V5 knock-in from figure (a) are shown, nuclear stain DAPI (blue) and TF-V5 fusion protein (yellow) are shown. (**c**) A 96-well plate heatmap of V5 knock-in for another set of 96 TFs. (**d**) Representative V5 knock-in panels, as obtained from the Operetta system, for the 96-well plate from (c) are shown, well number is indicated in the merged panel of each TF. Nuclear stain DAPI (blue), entire cell stain CellMask^TM^ HCS (green), and V5 tag (red) are shown. (**e**) Top 30 TFs with highest knock-in efficiency.

Remarkably, for the first 96-well plate, 30 out of 90 TFs were positive for V5 ICC with typical knock-in efficiencies ranging from 6%-29% (Figure 6A). A second 96 plate with distinct TFs showed similar knock-in efficiency (5%-38%), with 31 out of 95 TFs positive for V5 ICC (Figure 6C and 6D). These are similar efficiencies to those observed in our earlier single transfections (Figure 1 and 2). V5 ICC confirmed the expected nuclear localization of these TFs (Figure 6B). Thus, ~30% of genes were successfully tagged at the first attempt with good knock-in efficiency (Figure 6E).

*Cbfb* and *Ybx1*, showed V5 nucleocytoplasmic localization (Figure 6B and D). This illustrates the valuable information regarding protein localisation and levels data that can quickly obtained. The frequency of successful genes tagged in these experiments are likely to be an underestimate, as many TFs are likely low or non-expressed. These results clearly demonstrate the ease with which our method can be scaled.

### csRNP-derived knock-in lines can be used for immunoprecipitation-mass spectrometry (IP/MS) to identify protein partners

As a proof-of-principle of the applications we performed V5-immunoprecipitation followed by mass spectrometry (IP-MS), to identify interaction partners of *Olig2* in mouse GNS cells using the RIME (Mohammed et al. 2016) and ChIP-SICAP (Rafiee et al. 2016) methods. These enable identification of chromatin bound protein partners – the latter being more stringent for chromatin-bound Olig2. For each assay, we used V5 monoclonal antibody conjugated to magnetic beads. RIME analysis showed high enrichment of the bait protein Olig2 (Figure 7A). Subunits of SWI/SNF complexes and histone deacetylases (HDACs) in the pull-down complexes. Physical interaction of Olig2 and SWI/SNF complex has been previously reported and this interaction is essential for oligodendrocyte differentiation (Yu et al. 2013). HDACs have a known functional role in Olig2 function during development.

**Figure 7.**
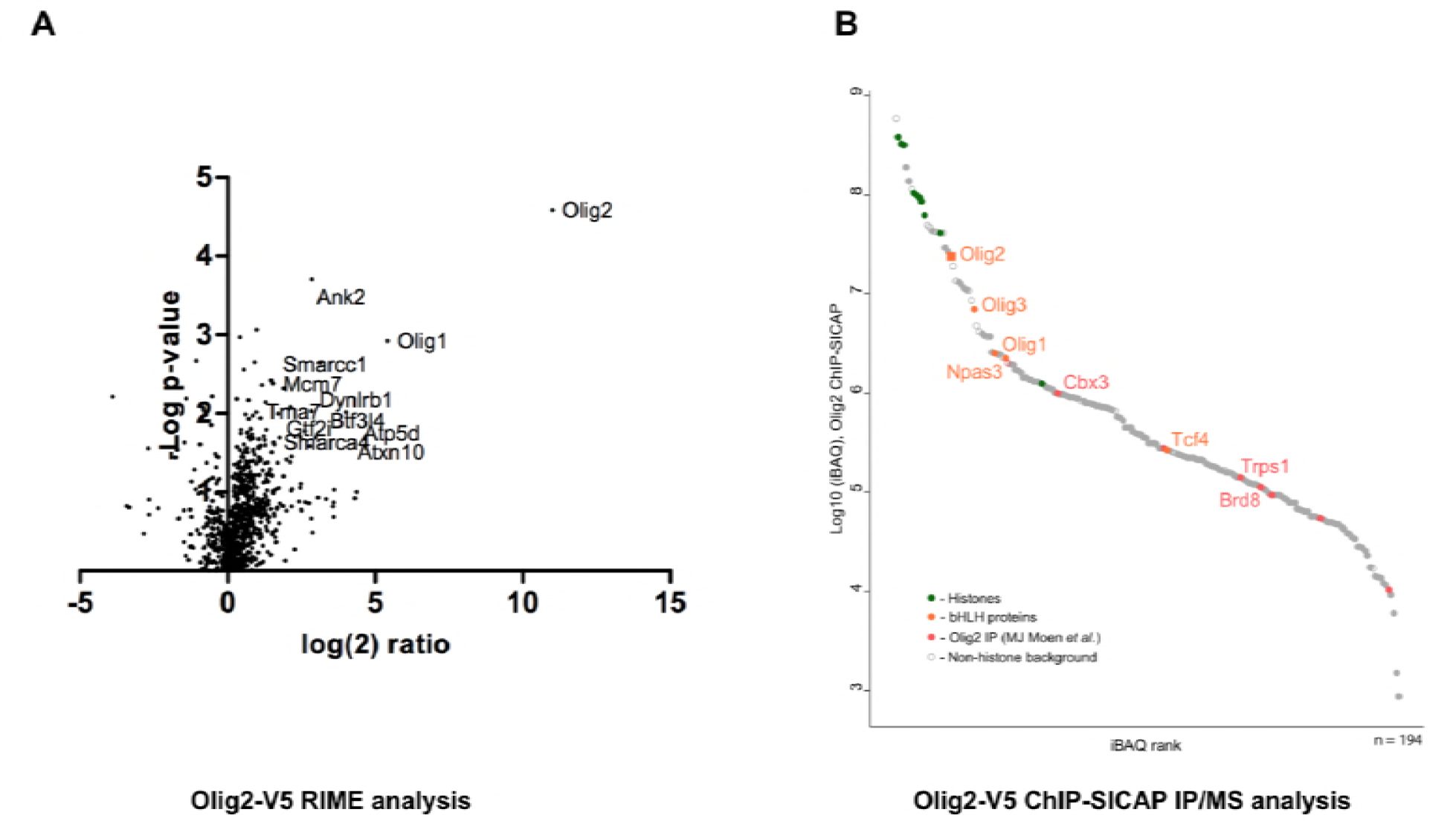
Identification of Olig2 partners using V5 knock-in lines. Mouse GNS Olig2-V5 cells were used for identification of Olig2 interaction partners. (a) RIME analysis, volcano plot showing log(2) fold change plotted against −log(10) p value for endogenously V5-tagged Olig2 samples versus samples generated from an untagged parental cell line (b) ChIP-SICAP analysis, proteins identified in the chromatin bound complexes are ranked based on iBAQ score in the descending order. (c) Olig2 interactions off chromatin from flow-through of (b).

ChIP-SICAP analysis showed strong enrichment of Olig2 and core histone proteins suggesting specific pull-down of chromatin fragments bound by Olig2 TF (Figure 7B). Noteworthy, two other oligodendrocyte lineage transcription factors, Olig1 and Olig3, were detected among the most enriched proteins. Earlier studies reported that sets of genes regulated by various Olig proteins have a partial overlap (Ligon et al. 2007; Meijer et al. 2012), explaining co-occupation of the same DNA-sites as shown by these data. Furthermore, we detected two other members of the basic helix-loop-helix (bHLH) family: Npas3 and Tcf4. The bHLH transcription factors are known to form heterodimers with other bHLH proteins on chromatin (Massari and Murre 2000) and the presence of Npas3 and Tcf4 may be explained by their direct physical interaction (Figure 7B, Supplementary Table 4). Both Npas3 and Tcf4 have been reported to be involved in CNS development (Shin and Kim 2013; Chen et al. 2016). Interestingly, a recent study reporting interactions of Olig2, Tcf4 and Npas3 in mouse neural stem cells by FLAG-affinity purification (Moen et al. 2017) allowed us to correlate interactomes of all the three bHLH proteins with our data. Taken together our results indicate that Tcf4 and Npas3 co-localize with Olig2 on chromatin, suggesting a functional interaction.

In addition to protein interactions of Olig2 on-chromatin, we also analyzed the flow-through of the streptavidin enrichment, representing interactions with soluble Olig2 (Supplementary Table 4). As expected, Olig2, Olig1, and Olig3 were found among the most highly abundant proteins, indicating high specificity of the immunoprecipitation. Furthermore, composition of identified proteins correlates with previous findings: we detected three known interactors of Olig2 (Cul3, Smarca4 and Sox8 (BioGRID, Intact)) and 55 other proteins reported to co-precipitate with Olig2, (Moen et al. 2017) (Fig 7). These included many SWI/SNF family members, and chromatin regulators Cbx3 and Chd4, consistent with RIME. Collectively, our data confirm that V5-tagging can be effectively combined with ChIP-SICAP to identify proteins that co-localize on chromatin or that interact off-chromatin.

## Discussion

An overarching goal in biology is to determine the key functions of each protein encoded in the genome. Epitope tagging of endogenous genes using CRISPR-assisted knock-in provides new opportunities to interrogate protein function, expression, subcellular localisation and interacting partners. In this study, we demonstrated simple and efficient epitope tag knock-in across a large number of genes in mouse and human primary stem cells.

Use of Cas9 recombinant protein is critical. We found that when combined with synthetic modified RNAs significant enhancement in the efficiency is possible. This was achieved using cheaper, faster and more reliable method – compared to plasmid based strategies. Importantly, we showed primary mammalian cells are readily amenable to RNP engineering, without the need for plasmid production, or prior genetic manipulation of the host cell lines. Our method is therefore versatile enough to be implemented by any laboratory in their existing cell lines. All reagents can now be obtained ‘off the shelf’.

Assembly of RNPs with csRNAs is simple and requires only ~30 min hands-on-time; this compared favourably to IVT reactions, which require multiple steps, are time-consuming, and result in variable quality of gRNAs. In our hands, these reagents are stable at 4oC for many months, and is particularly convenient when re-used across a range of cell types. Perhaps more importantly, use of the custom synthetic cr/tracrRNAs with their modified backbone and protected ends, shields the RNA from cellular RNases. We found greatly reduced toxicity – one of the key advantages of the RNP versus plasmids-based delivery.

We routinely generated clonal knock-in lines in a time-frame of 4-6 weeks. Consistent with a previous report (Liang et al. 2015), we found that RNP complexes target both copies of the genes at high frequency, enabling facile isolation of bi-allelic knock-in clones. Although monoallelic knock-in clones are sufficient for pull-down assays, bi-allelic clones are preferred to have more confidence in the interpretation of downstream assays.

While our focus has been on neural stem cells and their malignant counterparts (glioma stem cells), these same methods and reagents can work well in other stem cells, such as ES cells. Indeed, our knock-in data using ES cells revealed that non-expressed genes can be efficiently tagged using the same protocol. Under appropriate differentiation cues, induction of these proteins can be monitored using ICC analysis in the ES cell differentiating progeny. We believe this could be a key application – rapidly enabling assessment of proteins across a range of lineage contexts.

Despite the excitement, there are also some caveats. Foremost, inevitably there is a risk that C-terminus tagging can potentially compromise protein function, localisation or levels in the edited cells. Not all proteins will be amenable to tagging. If the protein of interest harbours critical C terminal domains, an N-terminus approach could be pursued – or knock-in to other regions such as structurally neutral linkers, if this is known.

Previous findings have reported a higher HDR rate with the guide RNAs that cut near the insertion site (Bialk et al. 2015; Paquet et al. 2016). Our results corroborate these findings and reveal that cut-to-insertion site distance is a critical factor for successful knock-in. The average size of CRISPR/Cas9-induced indels in mammalian cells has been reported to be around 1-5 bp (Paquet et al. 2016). We therefore recommend designing guide RNAs in the 3’UTR, preferably cutting 8-15 bp downstream of stop codon, to avoid interfering with the stop codon.

The ability to scale-up tagging to 96-well format required us to tackle the bottleneck in design. Our newly developed ‘Tag-IN’ tool simplifies gene sequence retrieval and designing crRNA and ssODN for medium-throughput experiments and is available from any web browser. This can be modified for use with alternative design parameters and distinct tags. We demonstrated one of the key applications by single-cell imaging of the levels and localisation for 60 different transcription factors in glioma stem cells. These were successfully tagged without any screening of gRNAs or ssODN; those that failed could be due to the gRNA does not working effectively or the ssODN donor being sub-optimal. These could be re-tested with replacements – particularly for the ssODN the other strand can be used and works effectively in many instances (data not shown). Alternatively, the protein maybe too low to detect, but might still be detected by PCR genotyping.

The pipeline that emerges is fully scalable and does not require sophisticated tools, expertise, or know-how. There are no significant bottlenecks in either the design, acquisition/production of reagents or delivery into cells. It was possible to generate such data with relatively little labour investment, and the whole pipeline from obtaining the reagents to imaging data was accomplished within three weeks.

Increasing the throughput further, we can envision systematic surveys of many hundreds or thousands of genes. Because tag-specific antibodies work universally across different cell types and species, this approach should allow cross-species comparisons and will also complement existing efforts to develop and characterise native antibodies. Furthermore, tagging of DNA-binding proteins will allow us to generate comprehensive genome-wide binding sites via ChIP-Seq for key transcription factors, chromatin-modifying enzymes, and other gene regulatory proteins. Although not a focus of the current study, we also find that csRNP is highly effective for gene knockout studies (data not shown) and others (Jacobi et al. 2017).

We demonstrated proof-of-principle in identification of Olig2 protein partners. This is likely a key application for future studies; simplifying and scaling up the ability to probe protein complexes in a range of cell types and cell states. Using the Olig2-V5 knockin lines we demonstrated on-chromatin partners in mouse glioma cells using optimised RIME and ChIP-SICAP methods, validating previously reported protein interaction partners. These optimised V5 RIME and ChIP-SICAP methods can now be deployed using the same optimised protocol for other mammalian proteins, particular for those with no good quality antibodies – a key advantage of epitope tagging.

With the remarkable developments in single-cell mRNA profiling, and plans for systematic RNA and protein atlases of mammalian cell types (Rozenblatt-Rosen et al. 2017), there is a greater need than ever to relieve the bottleneck of exploring protein products. Descriptive maps of cell types, while valuable, must be complemented by careful and detail molecular and cellular functional studies. Our findings suggest that epitope using CRISPR/Cas assisted knock-in is now simple and efficient enough that systematic annotation of many hundreds or thousands of endogenous will be possible in mouse and human primary cells.

## Experimental procedures

### Cell culture

Mouse and human NS and GNS cell lines were cultured essentially as described previously (Conti et al. 2005; Pollard et al. 2006). Laminin was purchased from Cultrex, R&D Systems. ANS4 and BL6 NS cell lines have been described previously. Mouse ES (Tg2a) cells were cultured in GMEM supplemented with 10% fetal calf serum, 1x non-essential amino acids,1x glutamine/sodium pyruvate, 1xLIF, 1x Pen/strep and 100μM of β-mercaptoethanol. Media was changed every day and cells passaged approximately every other day onto plates pre-coated with 0.1% gelatin. Differentiation of ES cell was performed as described previously (Pollard et al 2006) with 1x10^4^ cells per cm^2^ being seeded in N2B27 complete media for 7 days, with media being changed every 1-2 days.

MasterShef7 (MS7) human embryonic stem cells (hESCs) were rcultured in Essential 8 (E8) medium (Gibco, A1517001) on tissue culture plastic coated with Human Recombinant Laminin-521 (BioLamina, LN521) at 5μg/ml. Routine passaging was performed by incubating cells for 5 minutes at 37°C in 0.5 mM EDTA in PBS. Single-cell dissociation prior to nucleofection was performed by incubating cells for 10 minutes at 37°C in accutase. Y-27632 (Cell Guidance Systems, SM02) was included in the culture medium at 10 μM following initial thawing and after plating following nucleofection.

### Colony picking and clonal lines generation

Clonal cell lines were derived from the bulk populations using either single-cell deposition to 96-well plates or by manual colony picking. Single cells were deposited into 96-well plates using BD FACSAria™ II cell sorter. Depending on the cell lines, we obtained 30-40 colonies per 96-well plate in two weeks. For manual colony picking, mouse cells were seeded at clonal density (400 cells per 10 cm dish for NS cells, 100 cells per dish for GNS cells) to a 10cm dish and incubated in the complete media for 10-12 days for colony formation. From each dish, we picked 25-30 manually with a P200 pipette. Colonies from both methods were later replica plated into 96-well plates and analysed for successful knock-in using immunocytochemistry against the V5 tag. The V5 positive clones were further expanded for DNA extraction (PCR genotyping) and cryostorage.

### Design of guide RNAs and ssODN repair templates

For manual design: the 3’UTR sequence and 500bp sequence upstream of the 3’UTR were retrieved using Biomart tool. The final coding exon and 3’UTR features were manually annotated using SnapGene and the 200 bp around the stop codon were used as an input for guide RNA designing. We designed guide RNAs using either the web-based tool form Desktop Genetics (https://www.deskgen.com/landing/) or our own bioinformatics ‘Tag-IN’ tool (below). High scoring guide RNAs were picked for synthesis (i.e those with cut sites in the 3’UTR, preferably within 10-15 bp distance from the stop codon and no predicted off target cleavage). For ssODN design, first the PAM-blocking mutations (NGG>NGC or NGT) were introduced into the SnapGene sequence and then the epitope-tag coding sequence was inserted before the stop codon. The <200-mer ssODN ultramer was chosen to be the same strand as the guide RNA (also referred to as the PAM-strand, non-complementary strand or non-targeting strand) and is comprised of: a 5’ homology arm (~70-mer), the epitope tag coding sequence, stop codon, and a 3’ homology arm with the PAM-blocking mutations (~70-mer). For some of the ultramers the PAM-strand synthesis had failed and, therefore, the complementary strand (non-PAM strand) was synthesized as a donor DNA.

### Custom synthetic crRNA, tracrRNA, and ssODN

Custom synthetic crRNAs, tracrRNA, and ssODNs were manufactured by Integrated DNA Technologies, USA. The RNA backbone and ends were chemically modified for protection against cellular RNases. The 36-mer crRNA contains a variable gene-specific 20-nt target sequence followed by 16-nt sequence that base-pairs with the tracrRNA. The 67-mer tracrRNA contains the gRNA-scaffold sequence as well as 16-nt sequence complementary to crRNA. The lyophilized crRNA and tracrRNA pellets were resuspended in Duplex buffer (IDT) at 100 μM concentration and stored in small aliquots at -80 ⁰C. ssODN donor DNAs lyophilized pellets were supplied without modifications and resuspended in Duplex buffer (IDT) at 30 μM concentration.

### Production of *in vitro-transcribed* sgRNA

DNA template for T7-driven synthesis was prepared by annealing 119-mer, single-stranded, complementary ultramers (from IDT) encoding T7 promoter, guide RNA, and gRNA scaffold sequences. 200 ng of the template were used to synthesize sgRNA with the MEGAscript^®^ T7 Transcription Kit. The sgRNA was further purified using MEGAclear^TM^ Transcription Clean-Up Kit and stored at -80 ⁰C.

### Production of recombinant Cas9 protein

BL21(DE3) cells (New England Biolabs, C2527) were transformed with the plasmid pET28a/Cas9-Cys (Addgene, #53261) using standard protocols. Cas9 protein expression was induced with 0.5 mM IPTG (Isopropyl β-D-1-thiogalactopyranoside) (Fisher, 10715114) and the cells were incubated overnight at 20 ºC. 24hr later, bacterial pellets were resuspended in 20 ml of lysis buffer (20 mM Tris-HCl pH 8.0, 500 mM NaCl, 1mM TCEP, 5 mM imidazole pH 8.0), sonicated and loaded on a HisTrap HP 5 ml column (GE, 17-5248-01). The Cas9 protein was collected on elution buffer (20 mM Tris-HCl pH 8.0, 250 mM NaCl, 10% glycerol, 1 mM TCEP, 250 mM imidazole pH 8.0). The fractions containing Cas9 protein were pooled and loaded into a HiPrep 26/10 Desalting Column (GE, 28-4026-52) to equilibrate into Cas9 buffer (20 mM HEPES-KOH pH 7.5, 150 mM KCl, 1 mM TCEP). The purified Cas9 protein was further concentrated using Vivaspin columns (Vivaspin20, 30 000 MWCO PES, Sartorius stedim, VS2021) as per the users-guide instructions.

### Assembly of the active ribonucleoprotein (Cas9 plus csRNAs)

Synthetic Alt-R^®^ CRISPR/Cas9 crRNAs and tracrRNA were supplied by IDT. We prepared Cas9 RNP complexes immediately before electroporation experiments. Cas9 RNPs with IVT sgRNA were assembled (1-3 μg of IVT sgRNA with 5-10 μg of Cas9 protein) as described previously {Bressan:2017iz}. Briefly: 100 picomoles of each crRNA and tracrRNA were annealed using gradual step-down cooling in the PCR block (95 ⁰C at ramp rate 6 ⁰C/sec, 95 ⁰C - 25 ⁰C at ramp rate 0.1 ⁰C/sec, 25 ⁰C - 4 ⁰C (store) at ramp rate 0.5 ⁰C/sec).

Ribonucleoprotein (RNP) complexes were assembled by adding 10 μg of recombinant Cas9 protein to the annealed cr/tracrRNAs, incubated at room temperature for 10 minutes and stored on ice until electroporation into cells. 30 picomoles of single-stranded donor DNA were added to RNP complexes just before electroporation to prepare the complete RNP mix. For multiplex epitope tagging, 100 picomoles of cr/tracrRNA of each Sox2 and Olig2 were mixed together with 20 μg of rCas9 protein.

### Cell transfection

We used 4D Amaxa nucleofection system for the delivery of CRISPR ingredients. For NS cells and GNS cells, approximately 1 x 10^5^ cells were resuspended in 20 μL of Lonza SG cell line buffer and were mixed with the complete RNP mix and electroporated using the DN-100 program (two consecutive pulses for mouse cells) or using EN-138 program (one pulse for human cells). For embryonic stem cells, approximately 6 x 10^4^ cells in 20 μL of Lonza P3 primary cell buffer were used for each transfection with different programs: one pulse of program CA-120 for mouse ESCs; program CB-150 for human ESCs. After the electroporation, cells were transferred into a 6-well plate and allowed to recover for 3-5 days and later seeded into 96-well plates (1-2 x 10^4^ cells per well) for ICC.

For scale up, RNP assembly and delivery were performed as above, except that RNP complexes were prepared a day before and stored at −20 ⁰C. Electroporation was performed using the 96-well Shuttle^®^ device (Amaxa, Lonza). Immediately after transfection cells were transferred into a 96 deep-well plate and replica plates for immunocytochemistry assay were prepared by dispensing 1 x 10^4^ cells into 96-well plate using CyBi^®^-FeliX Liquid Handling Platform.

### Immunocytochemistry and imaging

We performed ICC on 96-well plates 5 days after transfection. Cells were washed once with PBS and fixed using 4% paraformaldehyde for 10min at room temperature and then permeabilized in PBST (PBS + 0.1% Triton X-100) for 20 min. Samples were incubated with blocking solution (1% goat serum in PBST) for 20 min at room temperature to block non-specific binding of the antibodies. Samples were treated overnight with primary antibodies in blocking solution followed by incubation with appropriate secondary antibodies and 4′,6-diamidino-2-phenylindole (DAPI). Images were acquired using either a Nikon wide-field fluorescence microscope or a PerkinElmer Operetta high-content imaging system. V5 positive cells were scored using Fiji software.

The following primary antibodies were used: V5 tag (eBioscience, TCM5 #14-6796-82,1:1000); HA Tag (Cell Signalling, 6E2 #2367, 1:100); FLAG tag (Sigma-Aldrich, #F3165, 1:2000); Myc tag (Cell Signalling, 9B11 #2276, 1:4000), Alexa Fluor secondary antibodies mostly Alexa Fluor Plus 647 (Thermo Fisher Scientific, 1:1000). HCS CellMask^TM^ Green Stain (Thermo Fisher Scientific, #H32714) for nucleocytoplasmic staining was used at 1: 10,000 for 20 min at room temperature.

### Genomic DNA extraction and PCR genotyping

Genomic DNA was extracted either using in-house lysis buffer as described previously (Raul’s paper, bulk populations from 96-well plate) or using DNeasy Blood & Tissue Kit (Qiagen, # 69506, for DNA extraction from clonal lines in a 24-well plate). PCR primers flanking the V5 tag were designed online using Primer3Plus to generate 400-600 bp amplicons. PCR genotyping and Sanger sequencing were done as described previously (reference). DNA samples were analysed using 2.5% agarose gels.

### ChIP-SICAP and RIME

Cells from seven T150 flasks (150 cm^2^) were cultured until 70-80% confluence and then dissociated into single-cells using accutase. To fix DNA-protein and protein-protein interactions, the cell pellet was resuspended in 1.5% methanol-free formaldehyde (Pierce) in 10 mL PBS for 10 min at room temperature. Excess formaldehyde was quenched by adding 125 mM glycine and incubated for 5 min at room temperature. Cells were washed twice with cold PBS and stored at -80 degrees until further use. ChIP-SICAP experiments were performed as described previously (Rafiee *et al.*, 2016). Briefly, chromatin from 20 million formaldehyde-fixed cells was sheared by sonication (Bioruptor Pico, Diagenode) down to 150 - 500 bp fragments, which were used as input for immunoprecipitation with anti V5 antibody (Abcam, 15828) overnight at 4°C. Antibody was captured with Protein-G beads (LIFE technologies, 10004D), the associated DNA was biotinylated by terminal deoxynucleotidyl transferase (Thermo Fisher, EP0162) in the presence of biotin-11-dCTP (Jena Bioscience, NU-809-BIOX). The antibody was eluted from the beads in 7.5% SDS with 200mM DTT and the released DNA-protein complexes were bound by streptavidin magnetic beads (NEB, S1420). After subsequent washes with SDS washing buffer (Tris-CL 1 mM, 1% SDS, 200 mM NaCl, 1 mM EDTA), 20% isopropanol and 40% acetonitrile, the beads were boiled in 0.1% SDS in 50 mM ammonium bicarbonate and 10 mM DTT at 95°C for 20 min. Proteins were digested overnight with trypsin at 37°C and the resulting peptides were purified with the SP3 protocol as previously described (Hughes *et al*., 2014) and analysed using an Orbitrap Fusion LC-MS system.

### crRNA/ssODN design tool

The implementation of our crRNA/ssODN design tool was completed in four stages: extraction of a target genomic sequence from GRCh38.p5 or GRCm38.p4 genome builds, retrieval of crRNA sequences matching the pattern N_20_NGG, scoring and ranking of each crRNA using the “Rule Set 2” (Doench et al. 2016) and “MIT” scoring models (Hsu et al. 2013), and design of each corresponding ssODN sequence.

To accommodate genomic sequence extraction, an SQL database of genomic coordinates was built using the Genomic Features package in R. This SQL database was used to retrieve coding DNA sequence (CDS) ranges upon user query with a desired Ensembl transcript Id. Given a CDS range, a genomic sequence was then extracted from a corresponding GRCh38.p5 or GRCm38.p4 Fasta file.

For each target genomic sequence, crRNAs were extracted limited to the pattern N_20_NGG. crRNAs were then ranked using two scoring models, “Rule Set 2” for assessing crRNA efficiency, and the MIT scoring system for crRNA specificity (Hsu et al. 2013; Doench et al. 2016). The former was utilized as a standalone script retrieved from the “sgRNA Designer” website (https://portals.broadinstitute.org/gpp/public/analysis-tools/sgrna-design) and the latter was implemented as documented on the “CRISPR Design” website (http://crispr.mit.edu/about). Off targets for each crRNA were found using the short read aligner tool Bowtie, searching up to 3 mismatches (Langmead et al. 2009). Off targets that then match the PAM pattern NAG and NGG were extracted from the Bowtie output. In addition, crRNAs that cut close to the stop codon (8-15 bp in the 3’UTR), and within the UTR region, were prioritized. Given a batch request, the top two ranking crRNAs were selected for output. A minimum distance of 30 bp from the stop codon was chosen as an additional threshold for batch processing.

To implement ssODN design, user-defined tags were inserted immediately 5^I^ proximal to the stop codon. PAM sequences were changed to minimise potential for Cas9 cleavage of donor sequences. Where the PAM sequence resided in the 3^I^ UTR our tool modified the NGG PAM to NGC. Intronic or exonic PAM changes instead aimed to produce silent mutations, and where this was not possible, aimed to reduce alterations in function by minimising differences in hydrophobicity and charge. Final ssODN sequences were limited to 200-mer including the tag sequence.

We therefore present the “*Tag-IN*” design tool, a novel crRNA and ssODN design tool aimed at streamlining CRISPR knock-in experimentation (www.stembio.crisprtag.ed.uk).

### Immunoprecipitation-mass spectrometry

The Olig2 protein interactors were identified using the Rapid immunoprecipitation mass spectrometry of endogenous protein (RIME) protocol. The nuclear fraction was resuspended using 1 ml of LB1 (50 mM HEPES KOH pH 7.5, 140 mM NaCl, 1mM EDTA, 10% glycerol, and 0.5% NP-40) with protease and phosphatase inhibitors. Lysate was cleared by centrifugation at 2000g for 5 min at 4°C and the pellet was resuspended with 1 ml of LB2 (10 mM Tris-HCL [pH 8.0], 200 mM NaCl, 1 mM EDTA, and 0.5 mM EGTA), and mixed at 4°C for 5 min. Lysate was cleared by centrifugation and the pellet was resuspended in 0.5 ml of LB3 (10 mM Tris-HCl [pH 8], 100 mM NaCl, 1 mM EDTA, 0.5 mM EGTA, 0.1% Na-deoxycholate, and 0.5% N-lauroylsarcosine). Samples were sonicated in a waterbath sonicator (Diagenode Bioruptor) and cleared by centrifugation. 10 µL V5 trap magnetic beads (MBL) were used per sample. IPs, washes and on-bead digests were performed using a Thermo Kingfisher Duo, all steps are at 5°C unless otherwise stated. Beads were transferred into 500μl of cleared lysate and incubated for 2 hours with mixing. Beads were then transferred for two washes in RIPA buffer and three washes in non-detergent lysis buffer. On-bead digest war performed by transferring the washed beads into 100μl 2M urea, 100 mM Tris-HCL pH 7.5, 1 mM DTT containing 0.3μg trypsin (Promega) per sample, beads were incubated at 27°C for 30 minutes with mixing to achieve limited proteolysis. The beads were then removed and tryptic digest of the released peptides was allowed to continue for 9 hours at 37°C. Reduced cysteine residues were alkylated by adding iodoacetamide solution to a final concentration of 50 mM and incubated 30 min at room temperature, in the dark. Trypsin activity was inhibited by acidification of samples to a concentration of 1% TFA. Samples were desalted on a C18 Stage tip and eluates were analysed by HPLC coupled to a Q-Exactive mass spectrometer as described previously (1). Peptides and proteins were identified and quantified with the MaxQuant software package (1.5.3.8), and label-free quantification was performed by MaxLFQ (2). The search included variable modifications for oxidation of methionine, protein N-terminal acetylation, and carbamidomethylation as fixed modification. The FDR, determined by searching a reverse database, was set at 0.01 for both peptides and proteins.

## Conflict of interest statement

AMJ and MAB are both employed by Integrated DNA Technologies (IDT), who sells reagents similar to some described herein. IDT is, however, not a publicly traded company and the authors do not own any shares or equity in IDT. No other authors have any financial interests or relationships with IDT; nor do they own any shares or equity.

**Table 1:** Summary of knock-in lines generated using RNP delivery. Table summarizing knock-in lines derived from mouse NS and GBM-model cells. The % bi-allelic knock-in among all V5 positive clones, as confirmed by PCR genotyping, is listed in the last column.

| Cell type | Gene | Total V5 positive clones |  | Bi-allelic knock-in clones |  |
| --- | --- | --- | --- | --- | --- |
| Mouse NS cells | Sox2 | 13/89 | 14.6% | 4/13 | 30.1% |
|  | Sox3 | 8/55 | 14.5% | 5/8 | 62.5% |
| Mouse GBM-model cells | Sox2 | 14/242 | 5.8% | 1/14 | 7.1% |
|  | Sox3 | 19/96 | 19.8% | 5/19 | 26.3% |
|  | Sox8 | 7/96 | 7.3% | 4/7 | 57.1% |
|  | Sox9 | 50/469 | 10.6% | 7/46 | 15.2% |
|  | FoxK2 | 2/96 | 2.1% | 1/2 | 50.0% |

Supplementary Table 1 | List of crRNA targets sequences

Supplementary Table 2 | List of primers for PCR genotyping

Supplementary Table 3 | List of primers for PCR genotyping

Supplementary Table 4 | Protein interactions of Olig2

## Acknowledgements

Vectors were provided by Addgene for Cas9 recombinant protein production (Addgene Plasmid 53261). ESCs were provided by Ian Chambers. hESC cultures were provided by Gillian Morrison following approval from the MRC UK Stem Cell Bank. Vivien for tissue culture, Bertrand Vernay provided support for microscopy and image analysis. We thank Donal O’Carroll, Ian Chambers, and Abdenour Soufi for helpful comments on the manuscript. The project was also supported by the BBSRC/EPSRC/MRC Synthetic Biological Research Centre (BB/M018040/1), part of the UK Research Councils’ investment in ‘Synthetic Biology for Growth’. S.M.P is a Cancer research UK Senior Research Fellow (A17368).

## References

Aida T, Chiyo K, Usami T, Ishikubo H, Imahashi R, Wada Y, Tanaka KF, Sakuma T, Yamamoto T, Tanaka K. 2015. Cloning-free CRISPR/Cas system facilitates functional cassette knock-in in mice. Genome Biol 16: 87.

Anderson EM, Haupt A, Schiel JA, Chou E, Machado HB, Strezoska Ž, Lenger S, McClelland S, Birmingham A, Vermeulen A et al 2015. Systematic analysis of CRISPR-Cas9 mismatch tolerance reveals low levels of off-target activity. J Biotechnol 211: 56-65.

Bhaya D, Davison M, Barrangou R. 2011. CRISPR-Cas systems in bacteria and archaea: versatile small RNAs for adaptive defense and regulation. Annu Rev Genet 45: 273-297.

Bialk P, Rivera-Torres N, Strouse B, Kmiec EB. 2015. Regulation of Gene Editing Activity Directed by Single-Stranded Oligonucleotides and CRISPR/Cas9 Systems. PLoS One 10: e0129308.

Bressan RB, Dewari PS, Kalantzaki M, Gangoso E, Matjusaitis M, Garcia-Diaz C, Blin C, Grant V, Bulstrode H, Gogolok S et al 2017. Efficient CRISPR/Cas9-assisted gene targeting enables rapid and precise genetic manipulation of mammalian neural stem cells. Development 144: 635-648.

Cameron P, Fuller CK, Donohoue PD, Jones BN, Thompson MS, Carter MM, Gradia S, Vidal B, Garner E, Slorach EM et al 2017. Mapping the genomic landscape of CRISPR-Cas9 cleavage. Nat Methods 14: 600-606.

Chen T, Wu Q, Zhang Y, Lu T, Yue W, Zhang D. 2016. Tcf4 Controls Neuronal Migration of the Cerebral Cortex through Regulation of Bmp7. Front Mol Neurosci 9: 94.

Cong L, Ran FA, Cox D, Lin S, Barretto R, Habib N, Hsu PD, Wu X, Jiang W, Marraffini LA et al 2013. Multiplex genome engineering using CRISPR/Cas systems. Science 339: 819-823.

Conti L, Pollard SM, Gorba T, Reitano E, Toselli M, Biella G, Sun Y, Sanzone S, Ying QL, Cattaneo E et al 2005. Niche-independent symmetrical self-renewal of a mammalian tissue stem cell. PLoS Biol 3: e283.

Dalvai M, Loehr J, Jacquet K, Huard CC, Roques C, Herst P, Côté J, Doyon Y. 2015. A Scalable Genome-Editing-Based Approach for Mapping Multiprotein Complexes in Human Cells. Cell Rep 13: 621-633.

Doench JG, Fusi N, Sullender M, Hegde M, Vaimberg EW, Donovan KF, Smith I, Tothova Z, Wilen C, Orchard R et al 2016. Optimized sgRNA design to maximize activity and minimize off-target effects of CRISPR-Cas9. Nat Biotechnol 34: 184-191.

Doudna JA, Charpentier E. 2014. Genome editing. The new frontier of genome engineering with CRISPR-Cas9. Science 346: 1258096.

Gavin AC, Aloy P, Grandi P, Krause R, Boesche M, Marzioch M, Rau C, Jensen LJ, Bastuck S, Dümpelfeld B et al 2006. Proteome survey reveals modularity of the yeast cell machinery. Nature 440: 631-636.

Hendel A, Bak RO, Clark JT, Kennedy AB, Ryan DE, Roy S, Steinfeld I, Lunstad BD, Kaiser RJ, Wilkens AB et al 2015. Chemically modified guide RNAs enhance CRISPR-Cas genome editing in human primary cells. Nat Biotechnol 33: 985-989.

Hsu PD, Scott DA, Weinstein JA, Ran FA, Konermann S, Agarwala V, Li Y, Fine EJ, Wu X, Shalem O et al 2013. DNA targeting specificity of RNA-guided Cas9 nucleases. Nat Biotechnol 31: 827-832.

Jacobi AM, Rettig GR, Turk R, Collingwood MA, Zeiner SA, Quadros RM, Harms DW, Bonthuis PJ, Gregg C, Ohtsuka M et al 2017. Simplified CRISPR tools for efficient genome editing and streamlined protocols for their delivery into mammalian cells and mouse zygotes. Methods 121-122: 16-28.

Jarvik JW, Telmer CA. 1998. Epitope tagging. Annu Rev Genet 32: 601-618.

Jinek M, Jiang F, Taylor DW, Sternberg SH, Kaya E, Ma E, Anders C, Hauer M, Zhou K, Lin S et al 2014. Structures of Cas9 endonucleases reveal RNA-mediated conformational activation. Science 343: 1247997.

Kelley ML, Strezoska Ž, He K, Vermeulen A, Smith A. 2016. Versatility of chemically synthesized guide RNAs for CRISPR-Cas9 genome editing. J Biotechnol 233: 74-83.

Kim S, Kim D, Cho SW, Kim J, Kim JS. 2014. Highly efficient RNA-guided genome editing in human cells via delivery of purified Cas9 ribonucleoproteins. Genome Res 24: 1012-1019.

Krogan NJ, Cagney G, Yu H, Zhong G, Guo X, Ignatchenko A, Li J, Pu S, Datta N, Tikuisis AP et al 2006. Global landscape of protein complexes in the yeast Saccharomyces cerevisiae. Nature 440: 637-643.

Langmead B, Trapnell C, Pop M, Salzberg SL. 2009. Ultrafast and memory-efficient alignment of short DNA sequences to the human genome. Genome Biol 10: R25.

Liang X, Potter J, Kumar S, Ravinder N, Chesnut JD. 2017. Enhanced CRISPR/Cas9-mediated precise genome editing by improved design and delivery of gRNA, Cas9 nuclease, and donor DNA. J Biotechnol 241: 136-146.

Liang X, Potter J, Kumar S, Zou Y, Quintanilla R, Sridharan M, Carte J, Chen W, Roark N, Ranganathan S et al 2015. Rapid and highly efficient mammalian cell engineering via Cas9 protein transfection. J Biotechnol 208: 44-53.

Ligon KL, Huillard E, Mehta S, Kesari S, Liu H, Alberta JA, Bachoo RM, Kane M, Louis DN, Depinho RA et al 2007. Olig2-regulated lineage-restricted pathway controls replication competence in neural stem cells and malignant glioma. Neuron 53: 503-517.

Lin S, Staahl BT, Alla RK, Doudna JA. 2014. Enhanced homology-directed human genome engineering by controlled timing of CRISPR/Cas9 delivery. Elife 3: e04766.

Ma H, Marti-Gutierrez N, Park SW, Wu J, Lee Y, Suzuki K, Koski A, Ji D, Hayama T, Ahmed R et al 2017. Correction of a pathogenic gene mutation in human embryos. Nature 548: 413-419.

Massari ME, Murre C. 2000. Helix-loop-helix proteins: regulators of transcription in eucaryotic organisms. Mol Cell Biol 20: 429-440.

Meijer DH, Kane MF, Mehta S, Liu H, Harrington E, Taylor CM, Stiles CD, Rowitch DH. 2012. Separated at birth? The functional and molecular divergence of OLIG1 and OLIG2. Nat Rev Neurosci 13: 819-831.

Merkle FT, Neuhausser WM, Santos D, Valen E, Gagnon JA, Maas K, Sandoe J, Schier AF, Eggan K. 2015. Efficient CRISPR-Cas9-mediated generation of knockin human pluripotent stem cells lacking undesired mutations at the targeted locus. Cell Rep 11: 875-883.

Mikuni T, Nishiyama J, Sun Y, Kamasawa N, Yasuda R. 2016. High-Throughput, High-Resolution Mapping of Protein Localization in Mammalian Brain by In Vivo Genome Editing. Cell 165: 1803-1817.

Moen MJ, Adams HH, Brandsma JH, Dekkers DH, Akinci U, Karkampouna S, Quevedo M, Kockx CE, Ozgür Z, van IJcken WF et al 2017. An interaction network of mental disorder proteins in neural stem cells. Transl Psychiatry 7: e1082.

Mohammed H, Taylor C, Brown GD, Papachristou EK, Carroll JS, D’Santos CS. 2016. Rapid immunoprecipitation mass spectrometry of endogenous proteins (RIME) for analysis of chromatin complexes. Nat Protoc 11: 316-326.

Paquet D, Kwart D, Chen A, Sproul A, Jacob S, Teo S, Olsen KM, Gregg A, Noggle S, Tessier-Lavigne M. 2016. Efficient introduction of specific homozygous and heterozygous mutations using CRISPR/Cas9. Nature 533: 125-129.

Pollard SM, Conti L, Sun Y, Goffredo D, Smith A. 2006. Adherent neural stem (NS) cells from fetal and adult forebrain. Cereb Cortex 16 Suppl 1: i112-120.

Rafiee MR, Girardot C, Sigismondo G, Krijgsveld J. 2016. Expanding the Circuitry of Pluripotency by Selective Isolation of Chromatin-Associated Proteins. Mol Cell 64: 624-635.

Rahdar M, McMahon MA, Prakash TP, Swayze EE, Bennett CF, Cleveland DW. 2015. Synthetic CRISPR RNA-Cas9-guided genome editing in human cells. Proc Natl Acad Sci U S A 112: E7110-7117.

Ramakrishna S, Kwaku Dad AB, Beloor J, Gopalappa R, Lee SK, Kim H. 2014. Gene disruption by cell-penetrating peptide-mediated delivery of Cas9 protein and guide RNA. Genome Res 24: 1020-1027.

Rivera-Torres N, Banas K, Bialk P, Bloh KM, Kmiec EB. 2017. Insertional Mutagenesis by CRISPR/Cas9 Ribonucleoprotein Gene Editing in Cells Targeted for Point Mutation Repair Directed by Short Single-Stranded DNA Oligonucleotides. PLoS One 12: e0169350.

Rozenblatt-Rosen O, Stubbington MJT, Regev A, Teichmann SA. 2017. The Human Cell Atlas: from vision to reality. Nature 550: 451-453.

Sander JD, Joung JK. 2014. CRISPR-Cas systems for editing, regulating and targeting genomes. Nat Biotechnol 32: 347-355.

Savic D, Partridge EC, Newberry KM, Smith SB, Meadows SK, Roberts BS, Mackiewicz M, Mendenhall EM, Myers RM. 2015. CETCh-seq: CRISPR epitope tagging ChIP-seq of DNA-binding proteins. Genome Res 25: 1581-1589.

Schumann K, Lin S, Boyer E, Simeonov DR, Subramaniam M, Gate RE, Haliburton GE, Ye CJ, Bluestone JA, Doudna JA et al 2015. Generation of knock-in primary human T cells using Cas9 ribonucleoproteins. Proc Natl Acad Sci U S A 112: 10437-10442.

Shin J, Kim J. 2013. Novel alternative splice variants of chicken NPAS3 are expressed in the developing central nervous system. Gene 530: 222-228.

Thul PJ, Åkesson L, Wiking M, Mahdessian D, Geladaki A, Ait Blal H, Alm T, Asplund A, Björk L, Breckels LM et al 2017. A subcellular map of the human proteome. Science 356.

Xiong X, Zhang Y, Yan J, Jain S, Chee S, Ren B, Zhao H. 2017. A Scalable Epitope Tagging Approach for High Throughput ChIP-Seq Analysis. ACS Synth Biol 6: 1034-1042.

Yu Y, Chen Y, Kim B, Wang H, Zhao C, He X, Liu L, Liu W, Wu LM, Mao M et al 2013. Olig2 targets chromatin remodelers to enhancers to initiate oligodendrocyte differentiation. Cell 152: 248-261.

Zuris JA, Thompson DB, Shu Y, Guilinger JP, Bessen JL, Hu JH, Maeder ML, Joung JK, Chen ZY, Liu DR. 2015. Cationic lipid-mediated delivery of proteins enables efficient protein-based genome editing in vitro and in vivo. Nat Biotechnol 33: 73-80.

